# GJA1-20k, an internally translated isoform of Connexin 43, is an actin capping protein

**DOI:** 10.1101/2022.01.05.475034

**Authors:** Rachel Baum, Joseph A. Palatinus, Miriam Waghalter, Daisuke Shimura, Qianru Jin, Lucas Kuzmanovich, Shaohua Xiao, André G. Kléber, Elena E. Grintsevich, TingTing Hong, Robin M. Shaw

**Affiliations:** Nora Eccles Harrison Cardiovascular Research and Training Institute, University of Utah, Salt Lake City, UT 84112, USA; Department of Biomedical Sciences, Cedars-Sinai Medical Center, Los Angeles, CA 90048, USA; Harvard Medical School Beth Israel and Deaconess Medical Center, Department of Pathology, Harvard University, Boston, MA 02134, USA; Department of Neurology, University of California, Los Angeles (UCLA), Los Angeles, CA 90095, USA; Department of Chemistry and Biochemistry, University of California, Los Angeles (UCLA), Los Angeles, CA 90095, USA; Department of Chemistry and Biochemistry, California State University, Long Beach (CSULB), Long Beach, CA 90840, USA; Diabetes and Metabolism Research Center, University of Utah, Salt Lake City, UT 84112, USA; Department of Pharmacology and Toxicology, College of Pharmacy, University of Utah, Salt Lake City, UT 84112, USA

## Abstract

Previously, we identified that GJA1-20k, an internally translated isoform of Connexin 43, mediates an actin-dependent protective form of mitochondrial fission (Shimura, Nuebel et al. 2021). We found that when GJA1-20k is present, bands of actin surround mitochondria at locations enriched with GJA1-20k, inducing mitochondrial fission which generates less oxygen free radicals, protecting hearts subjected to ischemia-reperfusion injury. Here, we report that GJA1-20k is a direct actin binding protein and thereby identify the mechanism by which GJA1-20k is able to recruit and stabilize actin filaments around the mitochondria. Surprisingly, GJA1-20k functions as a canonical actin capping protein, producing both truncated actin puncta and stabilized actin filaments. GJA1-20k contains an RPEL-like actin binding motif, and we confirm with both computational modeling and biochemistry, that this domain is crucial for actin capping. The actin capping functionality of GJA1-20k adds GJA1-20k to the family of proteins that regulate actin dynamics. As a stress responsive protein, GJA1-20k can help explain cytoskeletal dependent responses to cellular stress, from delivery of channels to affecting mitochondrial size and function.

## Introduction

Connexin 43 (Cx43) is a widely expressed gap junction protein that enables communication between cells. In the heart, rapid and targeted delivery of newly synthesized Cx43 protein to cell-cell borders is required to maintain gap junction plaques (Shaw, Fay et al. 2007, Xiao, Shimura et al. 2020) which function as low resistance pathways for ions to transit between adjacent cells, essential for the generation of each heartbeat. Cx43 is encoded by a single coding exon, *GJA1*, and therefore, its mRNA cannot be altered by splicing to generate multiple, slightly different, transcripts and isoforms. However, this single Cx43 mRNA is subject to internal translation, whereby internal AUG methionine sequences function as ribosomal start sites, generating N-terminus truncated isoforms (Smyth and Shaw 2013, Salat-Canela, Sesé et al. 2014).

GJA1-20k, the most abundantly expressed of up to six smaller isoforms (Smyth and Shaw 2013), is 20kDa in size and contains the Cx43 C-terminus tail but no full transmembrane domain. GJA1-20k is critical for full-length Cx43 to be trafficked to the cell border (Smyth and Shaw 2013, Xiao, Shimura et al. 2020). GJA1-20k also has channel independent roles in mitochondrial dynamics and biogenesis (Fu, Zhang et al. 2017, Basheer, Fu et al. 2018) as well as ischemic preconditioning (Basheer, Fu et al. 2018). GJA1-20k preferentially localizes to the mitochondria and is enriched at the outer membrane (Fu, Zhang et al. 2017).

We recently identified that GJA1-20k expression results in smaller sized mitochondria by stabilizing actin filaments which surround mitochondria and directly mediate mitochondrial scission events (Shimura, Nuebel et al. 2021). This form of GJA1-20k mediated mitochondrial fission is Dynamin Related Protein 1 (DRP1)-independent and results in a metabolic protective phenotype which reduces oxygen consumption and decreases reactive oxygen species (ROS) production. These findings uncovered a mechanism by which upregulation of GJA1-20k can mediate ischemic-preconditioning (Basheer, Fu et al. 2018) and reduce ischemic damage in the heart. While actin was found to be necessary for GJA1-20k mediated mitochondrial fission (Shimura, Nuebel et al. 2021), a direct interaction between GJA1-20k and actin has yet to be identified.

Mitochondrial fission inclusive, there are multiple lines of evidence suggesting a direct interaction between GJA1-20k and actin. Previous studies have identified that GJA1-20k expression does not alter the total amount of actin protein yet results in an increase in the number of F-actin filaments in cells and results in the stabilization of actin filaments (Basheer, Xiao et al. 2017). Actin is also a crucial component in the GJA1-20k mediated intracellular trafficking of Cx43 to the cell border (Smyth James, Vogan Jacob et al. 2012, Basheer, Xiao et al. 2017), as well as GJA1-20k mediated mitochondrial fission (Shimura, Nuebel et al. 2021). However, a direct binding interaction between GJA1-20k and actin has not been identified.

In this study, we identify a novel binding interaction between GJA1-20k and actin. Interestingly, we find that GJA1-20k has the properties of a canonical actin capping protein, directly reorganizing actin into both puncta and stabilized filaments. GJA1-20k contains an RPEL-like domain which is critical for actin interaction. These results not only identify GJA1-20k as a novel actin binding protein, but also reveal the mechanism by which GJA1-20k is able to recruit and stabilize actin around the mitochondria to induce a protective form of fission.

## Results

### GJA1-20k expression results in an actin puncta phenotype

To explore how GJA1-20k expression influences actin dynamics, we overexpressed GJA1-20k (Fig. 1A) in HeLa cells, which contain low levels of endogenous Cx43 and GJA1-20k (Shaw, Fay et al. 2007, Basheer, Xiao et al. 2017), and used a LifeAct plasmid to visualize the dynamic actin cytoskeleton. We detected that GJA1-20k expression, as previously observed, induced thickened actin filaments (Basheer, Xiao et al. 2017). However, we also discovered that GJA1-20k expression induces a population of subcortical actin that was reorganized into non-linear, circular puncta. These puncta are not present in GFP-transfected negative control cells (Fig. 1B).

**Figure 1.**
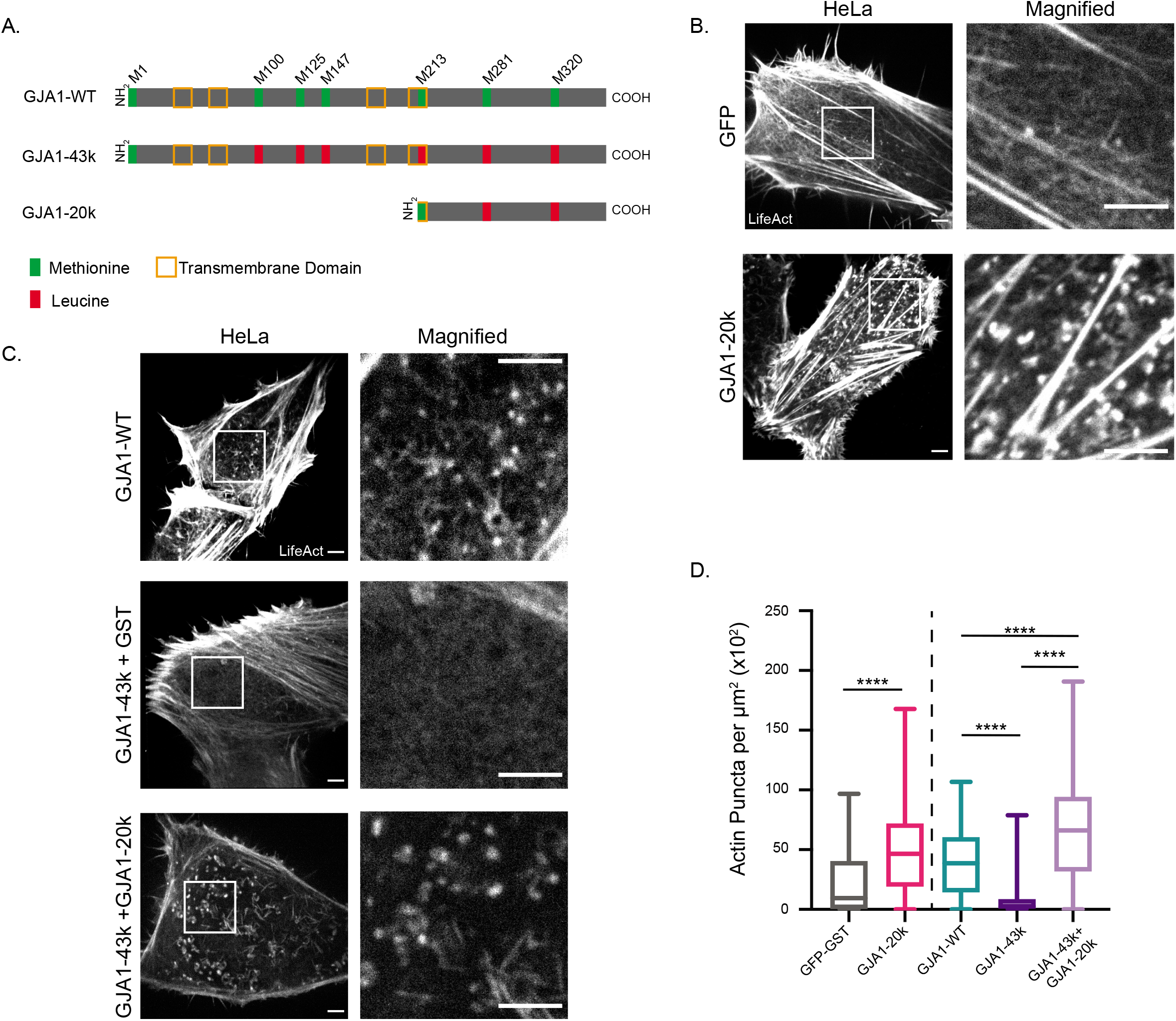
GJA1-20k expression results in a dual actin phenotype of actin puncta and stabilized actin filaments. A, Schematic of GJA1 plasmid designs used in overexpression experiments. Internal translation of truncated isoforms does not occur in plasmids where Methionines (green) are mutated to Leucines (red). B, Representative live cell confocal images of the actin cytoskeleton, visualized by LifeAct-mCherry, in HeLa cells expressing either 0.5μg of GJA1-20k-GFP or GST-GFP, scale bar=5μm. C, Representative live cell confocal images of the actin cytoskeleton, visualized by LifeAct-mCherry, in HeLa cells expressing either GJA1-WT (1μg), GJA1-43k-GFP(1μg) + GST-GFP(0.5μg), or GJA1-43k-GFP(1μg) + GJA1-20k-GFP(0.5μg), scale bar=5μm. D, Quantification of the total number of actin puncta per μm2 cell area of all HeLa transfection conditions. Data are presented as box and whisker plots with boxes showing median, 25th, and 75th percentile, with whiskers spanning the minimum to maximum range (n=100-200 cells per group) ****p<0.0001 by two sample t test (GFP-GST and GJA1-20k) and by one-way ANOVA with multiple comparisons with Bonferonni’s post-hoc test (GJA1-WT, GJA1-43k, and GJA1-43k + GJA1-20k).

We tested whether the appearance of actin puncta is specific to the GJA1-20k isoform of Cx43. HeLa cells were transfected with a wildtype Cx43 (GJA1-WT) plasmid that encodes full-length Cx43 and all downstream internally translated isoforms and compared to cells transfected with a full-length isoform that does not contain internal start sites (GJA1-43k) (schematic in Fig. 1A) and therefore does not produce any of the smaller isoforms. Actin puncta were present in cells expressing GJA1-WT but not GJA1-43k. The puncta in GJA1-43k treated cells were rescued with concurrent GJA1-20k expression (Fig. 1C), identifying GJA1-20k as a particular isoform generating the puncta. We generated a computational analysis algorithm that autonomously quantifies actin puncta in each cell. The algorithm found that GJA1-20k expression results in a significant increase in the number of actin puncta per cell area (Fig. 1D).

### Endogenous GJA1-20k expression results in the formation of actin puncta

To rule out off-target effects of exogenous expression, we explored whether the actin puncta can be generated by endogenously expressed GJA1-20k. C33A cells, which have high baseline levels of GJA1-20k (Salat-Canela, Sesé et al. 2014), were used. Immunoblotting confirmed that the relative levels of GJA1-20k in C33A cells were significant and exceeded the amount used in our HeLa cell overexpression experiments (Fig. 2A). Imaging using LifeAct to visualize actin revealed an actin puncta phenotype in C33A cells, similar to our GJA1-20k overexpression model (Fig. 2B). To confirm that endogenous GJA1-20k is responsible for the actin puncta phenotype, we used siRNA targeting *GJA1* to knock down expression of all GJA1 proteins and found a significant reduction in the number of actin puncta (Fig. 2C and D). Addition of a GFP negative control or full-length Cx43 did not rescue the actin puncta, however exogenous GJA1-20k expression did (Fig. 2C and D), further indicating that GJA1-20k expression is a specific isoform that can cause the actin puncta formation, whether expressed endogenously or exogenously.

**Figure 2.**
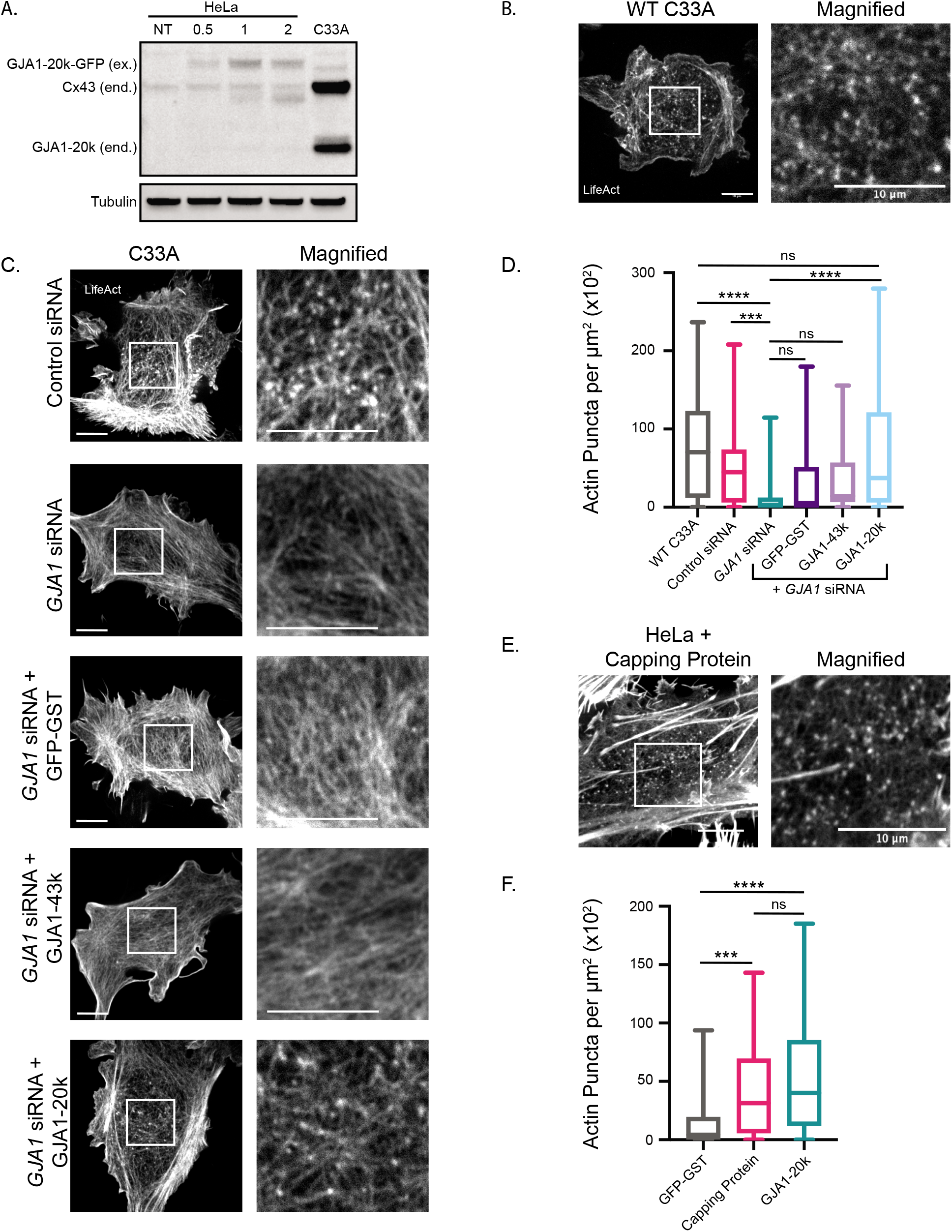
Endogenous GJA1-20k expression results in the formation of actin puncta. **A**, Western blot comparison of exogenous (ex.) GJA1-20k-GFP in HeLa cells and endogenous (end.) Cx43 and GJA1-20k in non-transfected C33A cells. HeLa cells were non-transfected (NT) or transfected with 0.5, 1, or 2μg of GJA1-20k-GFP. **B**, Representative live cell confocal imaging of actin, visualized by LifeAct-mCherry, in a wildtype C33A cell, or **C,** in C33A cells after siRNA treatment alone or with post siRNA knockdown expression of 0.5μg of either GFP-GST, GJA1-43k-GFP, or GJA1-20k-GFP, scale bars=10μm. **D**, Quantification of the total number of actin puncta per μm2 cell area for all C33A transfection cell conditions. Data are presented as box and whisker plots with boxes showing median, 25th, and 75th percentile, with whiskers spanning the minimum to maximum range (n=50-65 cells per group) ns=p>0.05, *p<0.05, **p<0.01, ***p<0.001, ****p<0.0001 by one-way ANOVA with multiple comparisons with Bonferonni’s post-hoc test. **E**, Representative live cell confocal LifeAct-mCherry tagged actin, in a HeLa cell expressing 0.5μg of Capping Protein plasmid, scale bars=10μm. **F**, Quantification of the total number of actin puncta per μm2 cell area for HeLa cells transfected with GFP-GST, Capping Protein, or GJA1-20k-GFP. Data are presented as box and whisker plots with boxes showing median, 25th, and 75th percentile, with whiskers spanning the minimum to maximum range (n=50-100 cells per group) ns=p>0.05, ***p<0.001, ****p<0.0001 by one-way ANOVA with multiple comparisons Bonferonni’s post-hoc test.

The formation of actin puncta (Fig. 1B-D, 2B-D) suggests that polymerization is being inhibited in the presence of GJA1-20k, which combined with our earlier findings that GJA1-20k stabilizes actin filaments (Basheer, Xiao et al. 2017, Shimura, Nuebel et al. 2021), led us to consider that GJA1-20k may be functioning as an actin capping protein. Capping proteins cap the barbed, or fast growing end of actin (Wear, Yamashita et al. 2003), preventing filament elongation by blocking the addition of new actin subunits while also stabilizing polymerized filaments by slowing barbed subunit dissociation (Kovar, Kuhn et al. 2003, Kueh, Brieher et al. 2008, Tang and Brieher 2013, Edwards, Zwolak et al. 2014). To compare phenotypes, we transfected HeLa cells with a plasmid encoding the actin interacting (Wear, Yamashita et al. 2003) alpha and beta subunits of Capping Protein (CP), a heterodimeric actin capping protein (Caldwell, Heiss et al. 1989, Cooper and Sept 2008), and observed a similar phenotype of actin puncta (Fig. 2E and F). Induction of cellular puncta is one indication that GJA1-20k and CP may interact with actin in a similar manner.

### GJA1-20k inhibits actin polymerization in a cell-free system

CP directly binds to actin (Wear, Yamashita et al. 2003). To explore whether GJA1-20k directly binds to actin and that it functions as a capping protein, we utilized total internal reflection microscopy (TIRFM) cell-free actin imaging, allowing us to visualize actin dynamics in the presence of purified proteins only. Purified CP was used as a positive control. We imaged actin after incubation across a range of CP concentrations. Consistent with previous studies of CP utilizing this assay (Huang, Blanchoin et al. 2003, Kovar, Kuhn et al. 2003, Dutta, Das et al. 2017), increased CP concentration results in decreased actin polymerization (Fig. 3A and B). Repeating the assay with purified GJA1-20k, we found that GJA1-20k, like CP, inhibits polymerization and results in a dose dependent decrease in actin polymerization (Fig. 3C and D), resulting in the puncta shape that we observed in our cell experiments.

**Figure 3.**
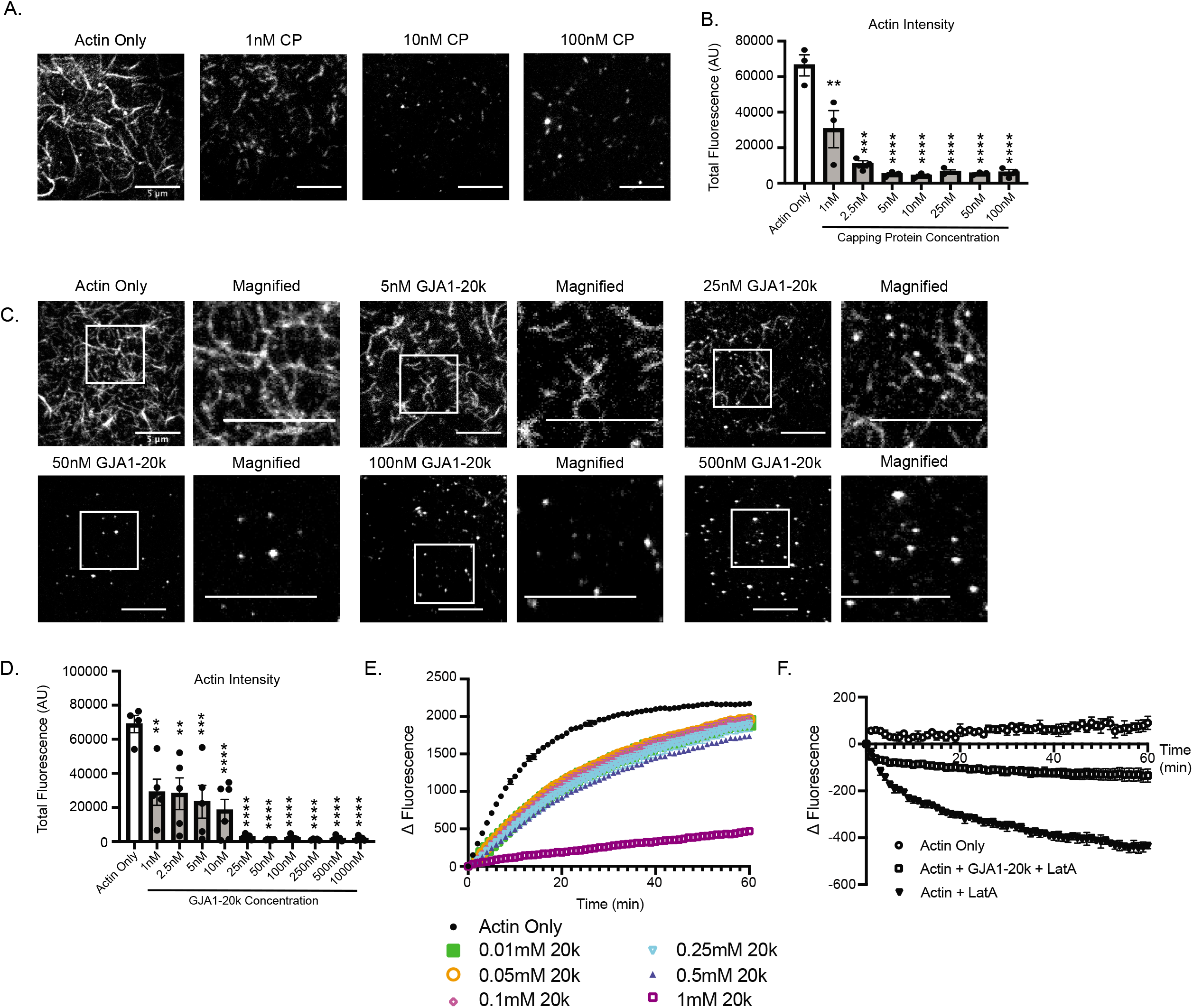
GJA1-20k inhibits actin polymerization. **A**, Representative TIRF images (t=15 minutes after initiation of polymerization) of cell-free Alexa Fluor 488 labeled actin in the presence of increasing concentrations of purified Capping Protein (alpha and beta subunits), scale bar=5μm. Reduction in actin polymerization is quantified by total fluorescence (AU) (**B**). Data are presented as mean±SEM (n=3 experiments per concentration) **p<0.01, ***p<0.001, ****p<0.0001 by one-way ANOVA with comparison to the Actin Only group with Bonferonni’s post-hoc test. **C**, Representative TIRF images (t=15 minutes after initiation of polymerization) of cell-free Alexa Fluor 488 labeled actin in the presence of increasing concentrations of GJA1-20k, scale bar=5μm. Reduction in actin polymerization is quantified by total fluorescence (AU) (**D**). Data are presented as mean±SEM (n=4-5 experiments per concentration) **p<0.01, ***p<0.001, ****p<0.0001 by one-way ANOVA with comparison to the Actin Only group with Bonferonni’s post-hoc test. **E**, Elongation of pyrene-labeled actin either alone or in the presence of purified GJA1-20k protein. Data are presented as mean±SD (n=3 separate experiments with duplicate wells for each group). **F**, Depolymerization of 0.1mM pyrene-labeled actin alone or in the presence of Latrunculin A or in the presence of Latrunculin A and purified GJA1-20k protein. Data are presented as mean±SD (n=3 separate experiments with duplicate wells for each group).

We previously reported that, in a cell-free actin polymerization assay, purified GJA1-20k stabilizes F-actin against depolymerization but has no significant effect on the inhibition of actin polymerization (Shimura, Nuebel et al. 2021). Based on our TIRF imaging findings (Fig. 3C and D), we wondered whether actin polymerization in a cell-free polymerization assay would be affected by increased concentrations of GJA1-20k. We found that increasing the concentration of GJA1-20k did inhibit actin polymerization in a dose-dependent manner (Fig. 3E), consistent with both our cell-free TIRF data and with the function of a capping protein. We also confirmed, using a cell-free actin depolymerization assay, that GJA1-20k stabilizes actin against depolymerization, also consistent with a capping protein phenotype (Fig. 3F).

### Structural modeling of the interaction between GJA1-20k and actin

Given direct interaction, we next sought to identify the actin binding domain on GJA1-20k. We could not identify homology between GJA1-20k and CP sequences. However, we were able to find a well-defined actin binding motif, the RPEL domain (Miralles, Posern et al. 2003), which has high sequence homology to the tail of GJA1-20k and binds to the same actin cleft as CP (Carlier, Pernier et al. 2015). The RPEL domain is defined as a sequence of RPxxxEL and it is conserved in known actin binding proteins such as MAL/MRTF-A myocardin-related transcription factor and the Phactr family of PP1-binding proteins (Guettler, Vartiainen et al. 2008, Mouilleron, Guettler et al. 2008, Mouilleron, Wiezlak et al. 2012). The last nine amino acids of GJA1-20k (RPRPDDLEI) (Fig. 4A) has a homologous domain, with two RP sequences and an Isoleucine substituted for the final Leucine in the RPxxxEL sequence.

**Figure 4.**
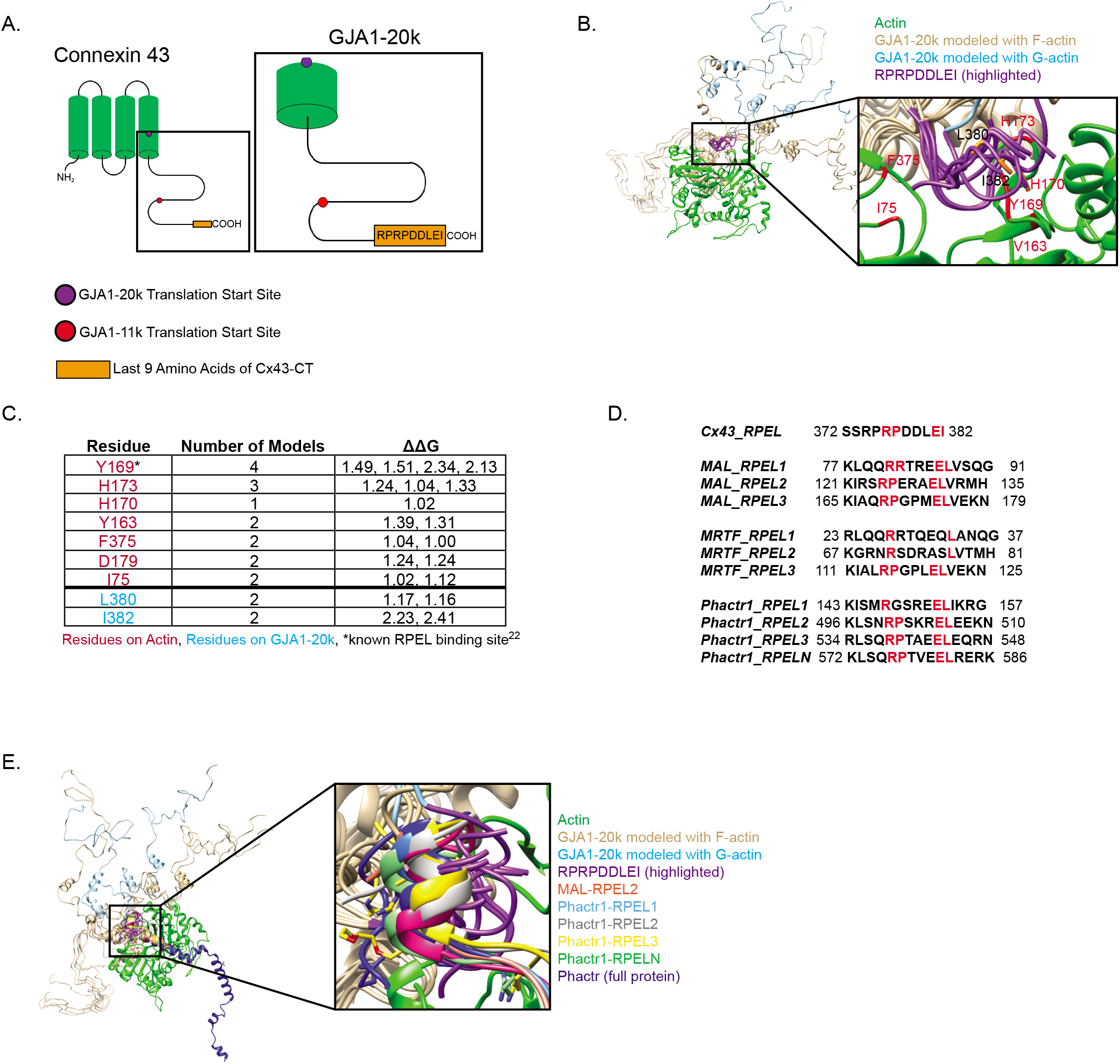
Structural Modeling of the Interaction between GJA1-20k and Actin. **A**, Illustration of Connexin 43 and GJA1-20k featuring the translation start sites for both the GJA1-20k and GJA1-11k isoforms and the sequence of the final nine amino acids (RPRPDDLEI). **B**, Modeling of GJA1-20k (PDB ID:1R5S) docked to actin. In tan: GJA1-20k ATTRACT models docked using F-actin (PDB ID:2ZWH), in light blue: GJA1-20k models docked using G-actin (PDB ID: 1J6Z), highlighted in purple: RPRPDDLEI tail sequence of GJA1-20k, red and black text in zoomed in box: hotspot residues as defined by Robetta Computational Alanine Scanning. Hotspots are identified by calculating the free energy change (ΔΔG>1) after a residue is mutated to alanine. **C**, Chart of hotspot residues on actin (red) and GJA1-20k (light blue) as defined by Robetta Computational Alanine Scanning. **D**, RPEL domain sequences of Cx43, MAL, MRTF-A, Phactr1. **E**, ATTRACT docking modeling of GJA1-20k (light blue and tan) compared to known RPEL domains of Phactr1 and MRTF-A. In orange: MAL-RPEL2, in pale blue: Phactr1-RPEL1, in silver: Phactr1-RPLE2, in yellow: Phactr1-RPEL3; in light green: Phactr1-RPELN, in dark blue: full Phactr1 protein.

To determine whether the sequence homology of GJA1-20k and the RPEL domain translates into predicted binding behavior, we used ATTRACT software (de Vries, Schindler et al. 2015) to model GJA1-20k docking on actin. Models of GJA1-20k were generated and, utilizing flexible protein-protein docking, we found that the tail sequence of GJA1-20k preferentially docks into the actin RPEL binding cleft (Fig. 4B). Each binding model was processed by Robetta Alanine Scanning, a computational alanine mutagenesis algorithm (Kortemme and Baker 2002, Kortemme, Kim et al. 2004), in order to quantify the stability of the interaction with actin. Seven hotspot residues on actin and two hotspot residues on GJA1-20k were identified (Fig. 4B and C) indicating sites which stabilize the protein-protein interaction. These results provide in silico evidence that the RPEL-like domain of GJA1-20k can theoretically bind in a stable fashion with actin.

While GJA1-20k does not have sequence homology with capping proteins, its RPEL-like domain may suffice as an actin binding domain. We compared the RPEL-like domain to the sequences (Fig. 4D) and structures (Fig. 4E) of known actin binding proteins that contain the RPEL domain, and confirmed homology. Our modeling identified that the tail of GJA1-20k aligns into the RPEL-binding actin cleft in a configuration similar to that of other RPEL proteins (Fig. 4E). Together, these modeling results suggest that GJA1-20k can potentially interact with actin via the RPEL-like domain in its terminal nine amino acids.

### The RPEL-like tail of GJA1-20k is sufficient to inhibit actin polymerization

We sought biochemical confirmation of our modeling data. The GJA1-20k sequence itself contains a downstream internal methionine which translates into a smaller Cx43 isoform, GJA1-11k (Smyth and Shaw 2013, Epifantseva, Xiao et al. 2020). Like GJA1-20k, GJA1-11k retains the RPEL-like C-terminus (Fig. 4A). GJA1-11k is a Cx43 isoform that localizes to the nucleus (Epifantseva, Xiao et al. 2020), so we investigated whether GJA1-11k expression alone affects actin localization. We found that GJA1-11k expression in HeLa cells resulted in an increase in nuclear actin retention (Fig. 5A-C), indicating that interaction with the RPEL-like domain organizes actin whether outside the nucleus (Figure 1B and C) or inside the nucleus (Figure 5A-C).

**Figure 5.**
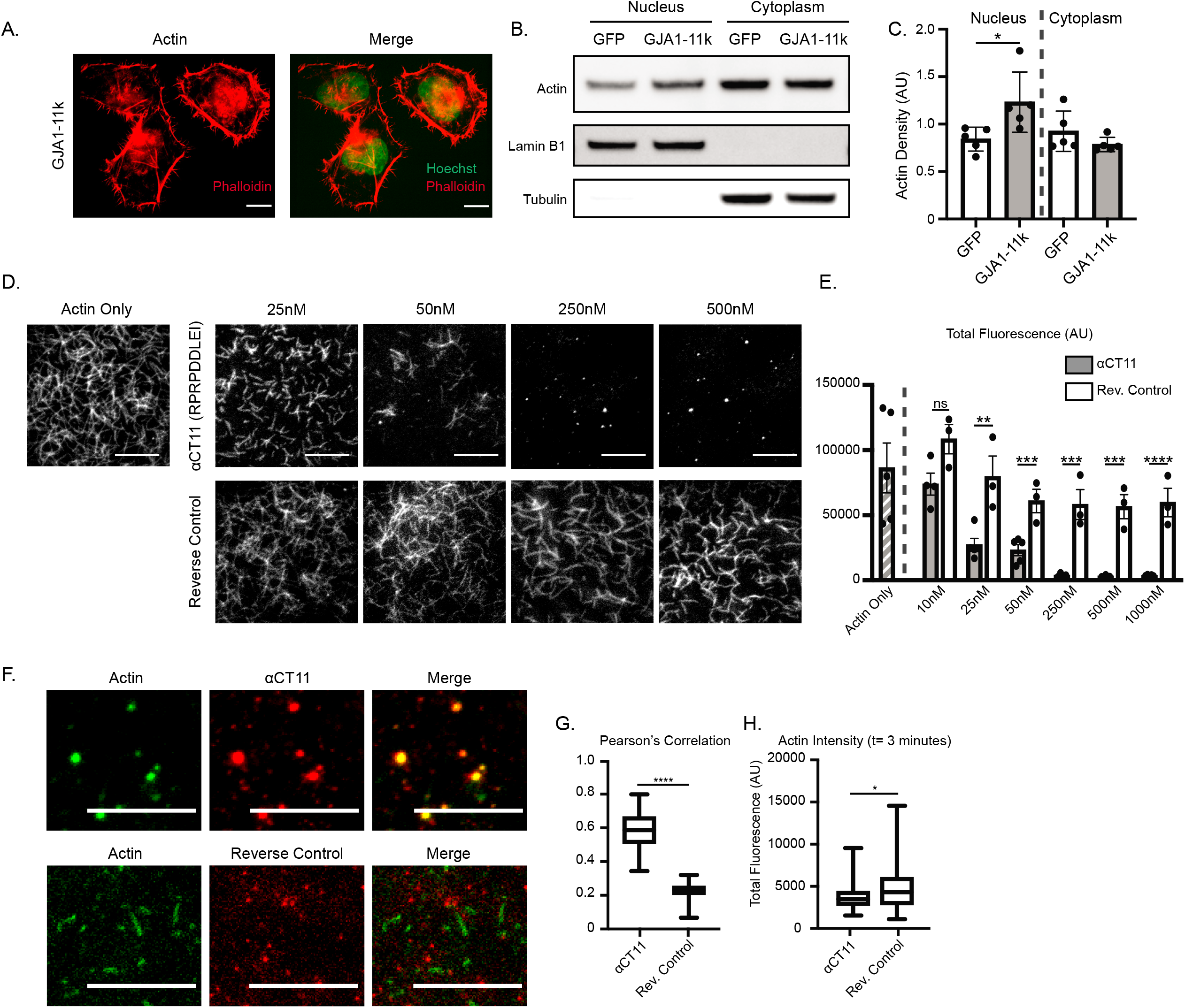
The RPEL-like tail of GJA1-20k is sufficient to inhibit actin polymerization. **A**, Representative immunofluorescence images of actin (phalloidin) and nuclei (Hoechst) in HeLa cells transfected with 0.5μg of GJA1-11k-V5, scale bar=10μm. **B**, Representative western blotting of actin in nuclear and cytoplasmic subcellular fractions of HeLa cells transfected with 0.5μg of either GJA1-11k-V5 or a GFP-V5 negative control. Lamin-B1 was used as a loading control for the nuclear fraction and Tubulin was used as a loading control for the cytoplasmic fraction. **C**, Quantification of western blots of actin density in the nuclear and cytoplasmic fraction of HeLa cells transfected with either GJA1-11k-V5 or GFP-V5, normalized to either Lamin-B1 (nucleus) or Tubulin (cytoplasm) loading controls. Data are presented as mean±SD (n=5 separate fractionation experiments) *p<0.05 by two sample t test. **D**, Representative TIRF images of cell-free Alexa Fluor 488 labeled actin in the presence of increasing concentrations of the αCT11 peptide (RPRPDDLEI) or a Reverse Control (IELDDPRPR), scale bar=5μm. Reduction in actin polymerization is quantified by total fluorescence (AU) (**E**). Data are presented as mean±SEM (n=3-5 experiments per concentration) ns=p>0.05, **p<0.01, ***p<0.001 ****p<0.0001 by multiple t tests comparing the αCT11 peptide to the Reverse Control for each concentration with Holm-Sidak correction for multiple comparisons. No significant difference was found between the Actin Only group and the Reverse Control groups (all concentrations), ns=p>0.05 by one-way ANOVA with multiple comparisons. **F**, Representative TIRF images of Alexa Fluor 488 labeled actin (green) in the presence of 250nM of Alexa Fluor 555 labeled αCT11 (red) or Reverse Control (red) to show colocalization (merge), scale bar=5μm, images taken 2-3 minutes after the initiation of polymerization. Images were quantified by using Pearson’s Correlation to measure colocalization (**G**) and by total fluorescence (AU) of the actin channel to measure changes in filament length (**H**). Data are presented as box and whisker plots with boxes showing median, 25th, and 75th percentile, with whiskers spanning the minimum to maximum range (n=35 frames from 3 separate experiments) *p<0.05, ****p<0.0001 by two sample t test.

To explore whether the RPEL-like sequence is sufficient to bind to actin, we utilized a peptide containing the last nine amino acids of GJA1-20k (RPRPDDLEI). This peptide, called αCT11, has been identified to be therapeutic in the setting of ischemia-reperfusion (Jiang, Hoagland et al. 2019) injury and as a mediator of actin reorganization (Chen, Mayo et al. 2015). We used the αCT11 peptide and a negative control containing the reverse sequence (IELDDPRPR) in the TIRFM cell-free actin assay and found that the αCT11 sequence alone was sufficient to cap actin filaments (Fig. 5D and E). Fluorescently conjugating each peptide for use in the TIRFM assay resulted in αCT11 colocalizing with actin at a significantly higher rate than the reverse control (Fig. 5F and G) and also significantly impaired actin filament growth (Fig. 5H), indicating that this domain of GJA1-20k retains the defining characteristic of capping proteins (Caldwell, Heiss et al. 1989).

### Functional similarities between GJA1-20k and capping proteins

Although we were able to confirm that the αCT11 sequence of GJA1-20k binds to and is sufficient to cap actin, capping proteins commonly have multiple actin interacting sites. For instance, CP has two binding sites that each bind to the barbed end of double-stranded actin (Wear, Yamashita et al. 2003) and the mutation or loss of either subunit decreases binding and capping affinity (Schafer, Hug et al. 1995, Hart and Cooper 1999, Kim, Yamashita et al. 2004). Other capping proteins, such as Gelsolin (Bryan 1988), Eps8 (Hertzog, Milanesi et al. 2010), and Villin (Khurana and George 2008), are also known to have multiple interaction sites with actin. To examine whether there is a second actin binding site on GJA1-20k, we deleted the αCT11 sequence from the C-terminus of GJA1-20k (GJA1-20kΔαCT11), fluorescently conjugated the protein, and performed the cell-free TIRFM assay to examine colocalization. We found that GJA1-20kΔαCT11 retains some colocalization with actin (Fig. 6A and B) suggesting a second binding site, yet without αCT11, the ability of GJA1-20kΔαCT11 to suppress actin polymerization is significantly diminished (Fig. 6C).

**Figure 6.**
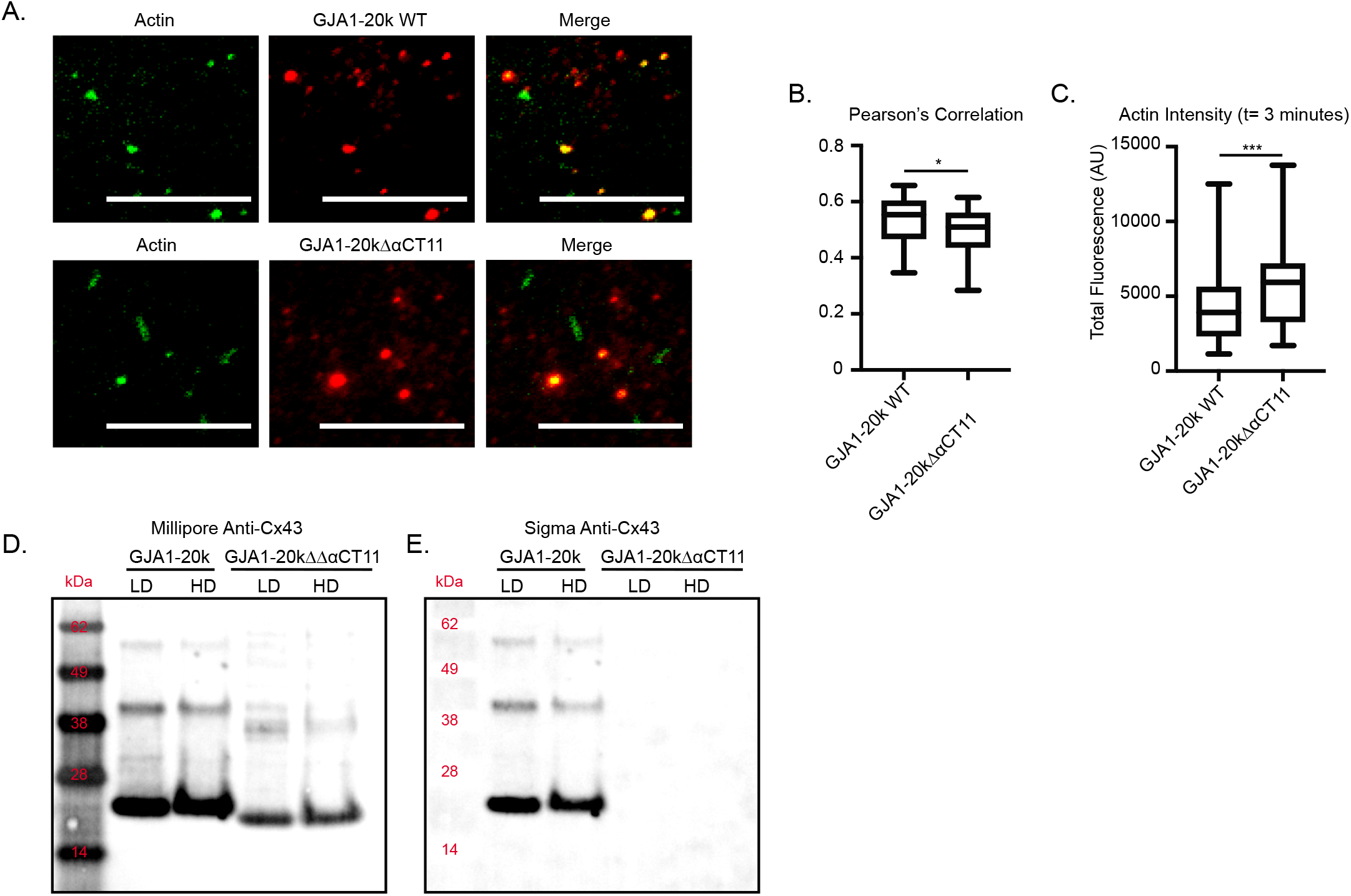
Characterizing GJA1-20k as an actin capping protein. **A**, Representative TIRF images of Alexa Fluor 488 labeled actin (green) in the presence of 250nM of Alexa Fluor 555 labeled GJA1-20k WT (red) or GJA1-20kΔαCT11 (red) to show colocalization (merge), scale bar=5μm, images taken between 2-3 minutes after the start of polymerization. Images were quantified by colocalization using Pearson’s Correlation to measure colocalization (**B**) and by total fluorescence (AU) of the actin channel to measure changes in filament length (**C**). Data are presented as box and whisker plots with boxes showing median, 25th, and 75th percentile, with whiskers spanning the minimum to maximum range (n=45 frames from 3 separate experiments) *p<0.05, ****p<0.0001 by two sample t test. **D**, Western Blot of purified GJA1-20k and GJA1-20kΔαCT11 protein incubated either in 1X anionic detergent (LD) or 4X anionic detergent (HD), detected with the Millipore manufactured anti-Cx43 antibody which targets an epitope that is present in both proteins. **E**, The same membrane probed with the Sigma manufactured Cx43 antibody which targets the C-terminal sequence that has been deleted in GJA1-20kΔαCT11.

We explored other phenotypic similarities between GJA1-20k and capping proteins. CP dimerizes, with heterodimers composed of its alpha and beta subunits both of which interact with actin (Wear, Yamashita et al. 2003). To better understand how GJA1-20k is able to cap actin in a manner similar to dimeric CP, we questioned whether GJA1-20k is able to oligomerize. Note, GJA1-20k is a smaller isoform of full length Cx43 which typically forms connexon hexamers (Shaw, Fay et al. 2007), and thus is known to oligomerize. After incubating purified GJA1-20k and GJA1-20kΔαCT11 proteins at room temperature to allow for self-binding, we subjected the proteins to either low or high concentrations of anionic detergent. We observed that when subjected to low anionic detergent, both GJA1-20k (20 kDa) and GJA1-20kΔαCT11 (19 kDa) can be detected by anti-Cx43 antibody at molecular weights indicative of dimers (38-40 kDa) and trimers (57-60 kDa), with an expected shift down in the smaller sized GJA1-20kΔαCT11 bands (Fig. 6D). Subjection to higher anionic detergent resulted in a reduction in the dimer and trimer band densities (Fig. 6D and E). To confirm that the bands at the dimer and trimer molecular weights were oligomerized GJA1-20k or GJA1-20kΔαCT11, we probed the same membrane with the a different (Sigma) anti-Cx43 antibody which targets an epitope within the αCT11 sequence. This antibody was able to detect all GJA1-20k bands but was unable to detect any GJA1-20kΔαCT11 bands (Fig. 6E), confirming that the dimer and trimer bands are specifically GJA1-20k or GJA1-20kΔαCT11. These results indicate that GJA1-20k is able to oligomerize, uncovering another similarity to capping proteins and likely a key component to how GJA1-20k is able to cap double-stranded actin filaments.

## Discussion

Using both intact cells and cell-free systems, we identify GJA1-20k as an intriguing member of the family of actin capping proteins. In addition to previously observed thickened actin filaments in cells expressing GJA1-20k (Basheer, Xiao et al. 2017), we now identify that GJA1-20k expression, whether endogenous or exogenous, directly inhibits actin polymerization which, in cells, reorganizes actin into truncated filaments that appear as puncta (Figure 3C and D). Interestingly, results from pyrene-actin assays indicate that GJA1-20k, while preventing actin polymerization, is also able to stabilize pre-formed actin filaments (Figure 3F). These results reconcile a dual actin phenotype of puncta and thickened filaments, indicating that GJA1-20k can differentially reorganize intracellular actin networks based on the relative availability of actin in monomeric G-actin pools (to be organized as puncta) versus filamentous F-actin fibers (to be stabilized). This dual actin phenotype in cells is typical of known actin capping proteins.

While a second actin binding site in GJA1-20k is yet to be determined, we were able to identify that the RPEL-like actin binding domain on GJA1-20k is critical for actin capping (Figure 5). Future studies are also needed to further understand how the actin modulating capacity of GJA1-20k is less active when part of full-length Cx43, a larger and multi-transmembrane domain protein which still retains the GJA1-20k actin binding domain. Without a lipophilic transmembrane domain, GJA1-20k can freely exist in the cytoplasm to interact with actin filaments and monomers. Freedom of movement may be sufficient to explain enhanced actin interaction with GJA1-20k, but this question should be explored more extensively.

The results of this study provide insight to the previously established role of GJA1-20k as a mediator of ischemic-preconditioning protection via its influence on mitochondrial dynamics (Shimura, Nuebel et al. 2021). By stabilizing actin around the mitochondria, GJA1-20k induces mitochondrial fission which results in decreased reactive oxygen species and results in organ protection. Identifying GJA1-20k as an actin capping protein answers a previously unknown question of how GJA1-20k is able to both recruit actin to surround the mitochondria and stabilize actin for fission induction.

The function of GJA1-20k as a capping protein also helps explain other roles of GJA1-20k. Previous studies had already identified two populations of actin that are crucial for Cx43 trafficking; 1) Cx43 connexon containing vesicles in transit to the cell border pause at actin rest stops (Smyth James, Vogan Jacob et al. 2012) and 2) stabilized actin filaments are necessary to pattern the microtubule delivery pathway (Basheer, Xiao et al. 2017). While GJA1-20k expression was associated with actin (Basheer, Xiao et al. 2017), it was unclear if GJA1-20k was a direct participant in the actin dynamics and reorganization. Our current finding that GJA1-20k caps actin filaments, suggests that GJA1-20k may be able to chaperone Cx43 to the cell border by dynamically remodeling the actin cytoskeleton to produce both the actin puncta that serve as vesicle rest stops and the stabilized actin filaments required for Cx43 trafficking. In this model, GJA1-20k would help pattern the cytoskeleton trafficking highways for subsequent delivery of full length Cx43 channel.

In myocytes, CP, is found at the Z-disc, functioning to cap the barbed end of thin filaments (Casella, Craig et al. 1987), and is also a key component of cardiac muscle structure (Hart and Cooper 1999). Gelsolin, a capping protein that, unlike CP and GJA1-20k, also severs actin filaments, has roles in cardiac hypertrophy (Hu, Ho et al. 2014) and stress-induced actin remodeling in cardiomyocytes (Patel, Zhabyeyev et al. 2018). While we have identified a role for GJA1-20k as an actin capping protein in trafficking and regulating mitochondrial shape and ischemic protection, the diverse functions of capping proteins highlight more potential roles to study of GJA1-20k as an actin regulator in the heart.

Actin capping proteins are also typically identified as regulators of cell migration and motility (Cunningham, Stossel et al. 1991, Frittoli, Matteoli et al. 2011, Sinnar, Antoku et al. 2014). Capping protein (CP) has been identified as a necessary component of cell migration, as it can be found at the leading edge of lamellipodia where it maintains pools of shorter filaments necessary to generate force (Akin and Mullins 2008). Cell motility is inhibited in the absence of CP (Sinnar, Antoku et al. 2014). As GJA1-20k functions as a capping protein, its effect on motility and migration may also be of interest to cancer research given that Cx43 has been widely studied in cancer, as both a tumor suppressor and as an oncogene (Bonacquisti and Nguyen 2019).

## Materials and Methods

### Cell Culture and Transfection

Low passage HeLa and C33A cell lines were grown at 37°C in a humidified atmosphere with 5% CO_2_ in Dulbecco’s Modified Eagle Medium (DMEM, Thermo Fisher Scientific) high glucose with 10% fetal bovine serum (FBS), nonessential amino acids (Thermo Fisher Scientific), sodium pyruvate (Thermo Fisher Scientific), and Mycozap-CL (Lonza). For transient transfection experiments, cells were plated 24 hours prior to transfection and the transfection was carried out using FuGENE® HD (Promega) or Lipofectamine 2000 (Thermo Fisher Scientific) per the manufacturer’s protocol.

### RNA interference

RNA knockdown was performed as previously described (Epifantseva, Xiao et al. 2020). We utilized siRNA (Thermo Fisher Scientific) synthesized to knockdown human *GJA1* mRNA (sequence 5’ to 3’: GGGAGAUGAGCAGUCUGCCUUUCGU; cat. # HSS178257) and Stealth™ RNAi (Thermo Fisher scientific, cat. # 12935112) was used as a negative control. C33A cells were transfected with 100nM of either siRNA using Lipofectamine™ RNAiMAX (Thermo Fisher Scientific) and further transfections with siRNA resistant plasmids were performed 24 hours after siRNA treatment using FuGENE® HD (Promega).

### Live cell imaging

Live cell imaging was performed using a Nikon Eclipse T*i* microscope with an attached incubator using a 100x/0.75 Plan Apo objective and a spinning-disk confocal unit (Yokogawa) controlled by NIS Elements software. Cells were plated in 35mm glass bottom dishes (MatTek) and transfection and imaging occurred in the same dishes. For imaging, cell media was exchanged for pre-warmed Hanks’ Balanced Salt solution (HBSS) with 10% FBS.

To visualize the cytoskeleton, cells were transfected with a LifeAct-mCherry plasmid, which was generated as previously described(Smyth James, Vogan Jacob et al. 2012) using Gateway cloning technology (Invitrogen). Briefly, the LifeAct Entry clone was inserted into the pDEST-mCherry-N1 destination vector to generate the LifeAct peptide with a C-terminus mCherry tag. The GJA1 plasmids used were also generated as previously described (Smyth and Shaw 2013, Basheer, Xiao et al. 2017) using Gateway cloning technology. Briefly, entry clones containing human *GJA1* encoding full-length Cx43 and the GJA1-43k and GJA1-20k isoforms were inserted into C-terminal GFP and YFP destination vectors. GJA1-11k was inserted into a C-terminal V5 destination vector as previously described (Epifantseva, Xiao et al. 2020). The Capping Protein (F-actin-capping protein subunit alpha-1 and F-actin-capping protein subunit beta) plasmid was a gift from Antoine Jégou & Guillaume Romet-Lemonne (Addgene plasmid #89950).

### Pyrene-actin assays

The pyrene actin polymerization and depolymerization assays were performed using the Actin Polymerization Biochem Kit (Cytoskeleton) per the manufacturer’s protocol with modifications. For the polymerization assay, pyrene labeled actin (Cytoskeleton) was reconstituted in G-buffer (5mM Tris-HCl, pH 8.0, 0.2mM CaCl_2_, 0.2mM ATP) to a final concentration of 2μM, and was then incubated on ice for 1 hour to depolymerize any F-actin, and then centrifuged at 14k rpm for 30 minutes at 4°C to remove any polymerized filaments. Actin was then polymerized either alone or in the presence of GJA1-20k (final concentration 0.1mM) in 0.25x polymerization buffer (Cytoskeleton). For the depolymerization assay, actin was prepared at a final concentration of 2μM in G-buffer and polymerized in 0.25x actin polymerization buffer at RT for 1 hour. Actin was maintained at a final working concentration of 2μM in F-buffer (G-buffer + 0.25x polymerization buffer). Actin was incubated with or without GJA1-20k (0.1mM) before depolymerization. Depolymerization was induced using 10μM Latrunculin A (Sigma-Aldrich) in DMSO, and an equal volume of DMSO was added to the Actin Only group. Pyrene fluorescence was measured using a SpectraMax M5e microplate reader with excitation of 355 nm and emission of 405nm at RT.

### Western Blotting

Western blotting was performed as previously described (Xiao, Shimura et al. 2020). Briefly, cell lysates were subjected to SDS-PAGE electrophoresis using NuPAGE 4-12% Bis-Tris gels and MES running buffer (Thermo Fisher Scientific) according the manufacturer’s protocol. Gels were then transferred to FluoroTrans PVDF Membranes (Pall). For antibody staining, membranes were blocked at room temperature (RT) in 5% nonfat dry milk (Carnation) in TNT buffer (50 mM Tris pH 8.0, 150 mM NaCl, 0.1% Tween-20) and probed with primary antibodies in blocking buffer at 4°C overnight. Membranes were then incubated with host-matched fluorophore-labeled secondary antibodies (1:1000, Invitrogen) at RT for 1 hour. Western blots were visualized using the ChemiDoc MP System and band density was quantified using ImageJ.

To detect endogenous and exogenous GJA1-20k (Fig. 2A), we used the following antibodies diluted in TNT buffer: anti-Cx43 (1:1000, Millipore) and anti-Tubulin (1:5000, abcam).

### Cell Fractionation

For the experiments that required nuclear fractionation, we used the NE-PER™ Nuclear and Cytoplasmic Extraction Kit (Thermo Fisher Scientific) per the manufacturer’s protocol. Briefly, HeLa cells were transfected with either GJA1-11k-V5 or a GFP-V5 control and 24 hours after transfection they were harvested with Trypsin-EDTA and pelleted. The nuclear and cytoplasmic fractions were isolated by following the manufacturer’s protocol and using the proprietary extraction reagents. The subcellular fractionation lysates were analyzed via Western Blot. We used the following primary antibodies diluted in TNT buffer: anti-beta actin (1:1000, GenScript), anti-Lamin B1 (1:1000, abcam), anti-Tubulin (1:5000, abcam).

### TIRFM Cell-free Actin Assays

Total internal reflection fluorescence microscopy (TIRFM) was conducted using a Nikon Eclipse T*i* microscope with a TIRF Apo 100x objective and recorded with an iXon Ultra 897 EMCCD Camera (Andor) controlled by NIS Elements Software. Purified His-tagged GJA1-20k protein, Capping Protein, αCT11 peptide (RPRPDDLEI), or Reverse Control (IELDDPRPR), at specified concentrations, were diluted in 1x TIRF Buffer (2x TIRF: 20mM imidazole, pH 7.4, 100 mM KCl, 2 mM MgCl_2_, 2mM EGTA, 0.4 mM ATP (Cytoskeleton, Inc.), 100 mM DTT, 30 mM glucose, 40 μg/mL catalase, 200 μg/mL glucose oxidase, and 1% methylcellulose). Alexa-488 conjugated rabbit skeletal muscle G-actin (Life Technologies) was added to start the assay at a final concentration of 1μM and the mixture was added to flow chambers that were preincubated with *N*-Ethylmaleimide (NEM)-Heavy-Mero-Myosin II (Cytoskeleton, Inc.) and washed with 1% bovine serum albumin (Thermo Fisher Scientific) in High-salt TBS (2x HS-TBS: 100mM Tris-HCl, pH 7.6, 1.2M NaCl) and Low-salt TBS (2x LS-TBS: 100mM Tris-HCl, pH 7.6, 100mM NaCL). Once added to the flow chambers, imaging was immediately started with images captured every 10 seconds for 15 minutes.

Total fluorescence was calculated using ImageJ. Thresholding was used to highlight actin and remove background fluorescence. Integrated density (IntDen) was then measured and recorded for each sample.

### Protein Purification

GJA1-20k, Capping Protein, and GJA1-20kΔαCT11 were purified for the TIRFM Microscopy Assays. His-tagged GJA1-20k plasmid without the putative transmembrane region (amino acids 236-382 of the full-length human Cx43, NCBI reference NP 000156,1) was generated and purified. To create the plasmid, a 6 x His tag and linker was fused to the N-terminus of the GJA1-20k sequence and cloned into the pET301/CT-DEST vector via Gateway cloning (Invitrogen). The primers used to amplify the sequence are (5’ to 3’) GGGGACAAGTTTGTACAAAA AAGCAGGCTTCAGGAGGTATACATATGCATCATCATCATCATCACGGTGGTGGCG GTTCAGGCGGAGGTGGCTCTGTTAAGGATCGGGTTAAGGGAAAG (sense) and GGGGACCACTTTGTACAAGAAAGCTGGGTCTTACTAATCGTCATCATCGTCATCATCGTCATCATCACTTCCACCACTTCCACCGATCTCCAGGTCATCAGGCCG (antisense). The protein was expressed in LOBSTR (Kerafast) E.coli competent cells transformed with pTf16 chaperone protein tag (TaKaRa Bio) per the manufacturer’s protocol.

A His-tagged mouse Capping Protein (F-actin-capping protein subunit alpha-1 and F-actin-capping protein subunit beta) plasmid was obtained from Addgene as described above (Plasmid #89950). To generate GJA1-20kΔαCT11, the plasmid encoding His-tagged GJA1-20k, was mutated to delete the last nine amino acids (RPRPDDLEI), using the QuikChange Lightning Site-Directed Mutagenesis Kit (Agilent) as described in the mutagenesis section below. These proteins were expressed in One Shot™ BL21 (DE3) competent cells (Invitrogen).

All proteins were expressed in E. coli as specified above and protein expression was induced at 37°C with 1mM isopropyl-1-thio-β-galactopyranoside. The induced proteins were isolated from bacterial pellets lysed in B-PER Bacterial Protein Extraction Reagent (Thermo Fisher Scientific) containing cOmplete ULTRA Protease Inhibitor Tablets (Sigma Aldrich) using the HisPur Cobalt Purification Kit (Thermo Fisher Scientific) per the manufacturer’s protocol with buffer substitutions (300 mM NaCl, 50 mM NaH_2_PO_4_, 5 mM 2-mercaptoethanol, pH 8.0 for column equilibration and washing with the addition of 10 mM and 20 mM Imidazole for sequential washing steps and 300 mM NaCl, 50 mM NaH_2_PO_4_, 150 mM Imidazole, pH 8.0, 10% glycerol for elution). Purified proteins were concentrated and subjected to a buffer exchange into a final buffer containing 50 mM NaH_2_PO_4_, 150 mM NaCl, pH 8.0, 10% glycerol.

Biotin-tagged peptides of the last nine amino acids of GJA1-20k (αCT11, RPRPDDLEI) and a reverse control (IELDDPRPR) that were also used in the TIRFM Microscopy Assay were synthesized by Anaspec.

### Mutagenesis

Mutations to delete the αCT11 region (RPRPDDLEI) of GJA1-20k and GJA1-WT was performed using the QuikChange Lightning Site-Directed Mutagenesis Kit (Agilent) according to the manufacturer’s instructions.

To mutate GJA1-20k for GJA1-20kΔ αCT11 protein production, a plasmid containing the sequence described above was mutated using the primers (5’ to 3’) GTCGTGCCAGCAGCGGTGGAAGTGGTGG (sense) and CCACCACTTCCACCGCTGCTGGCACGAC (antisense).

To mutate GJA1-WT to create Cx43ΔαCT11 to study trafficking, the GJA1-WT-YFP plasmid was altered using the primers CAGTCGTGCCAGCAGCAAGCTTCGAATTCTGC (sense) and GCAGAATTCGAAGCTTGCTGCTGGCACGACTG (antisense).

### TIRFM Cell-free Actin Assay with Fluorescently Conjugated Proteins

For TIRFM experiments that required fluorescent conjugation, the proteins or peptides (GJA1-20k, GJA1-20kΔαCT11, αCT11, IELDDPRPR) were fluorescently conjugated using the Alexa Fluor 555 Microscale Protein Labeling Kit (Thermo Fisher Scientific) per the manufacturer’s protocol.

The fluorescently conjugated proteins were used in the TIRFM cell-free actin assay at a concentration of 250nM. The TIRFM cell-free actin assay was performed as described above and 15 images on different areas of the slide were captured within 2-3 minutes after the start of polymerization.

Quantification of colocalization was performed with ImageJ, using the JACoP (Just Another Colocalization Plugin) plugin.

### Quantification of Actin Puncta per Cell Area

We generated a novel image analysis macro to quantify the number of actin puncta per cell area. Actin was imaged in HeLa or C33A cells, using a LifeAct plasmid, via live cell confocal imaging and then each condition was blinded for analysis. Using ImageJ for the analysis, each cell was selected and individually outlined, within the cortical actin border and the outside was cleared. A macro was then run to subtract the background using a rolling ball radius of 5 pixels, the actin puncta were highlighted using an auto threshold to highlight the subcortical actin. The analyze particles function was then run with a size exclusion criterion of 0.03-0.58 microns and a circularity setting of 0.4-1.0 to exclude actin filaments and only detect circular actin puncta. The analyze particles count was then normalized to the area of each cell.

### Immunofluorescence Staining

To image actin in the nucleus, cells were transfected with either GJA1-11k-V5 or GFP-V5 and fixed in 4% PFA for 20 minutes at RT, 24 hours after transfection. The fixed cells were then permeabilized in 0.1% Triton in PBS and blocked in 5% goat serum for 1 hour at RT. The cells were then incubated in Phalloidin (1:1000, Thermo Fisher Scientific). The nuclei were then stained with Hoechst 3342 (1:2000, Thermo Fisher Scientific) for 20 minutes at RT.

Images were taken using the Nikon Eclipse T*i* microscope with a 100x/0.75 Plan Apo objective, a spinning disk confocal unit (Yokogawa), controlled by NIS Elements software.

### Protein Oligomerization

Purified GJA1-20k or GJA1-20kΔαCT11, stored in 50 mM NaH_2_PO_4_, 150 mM NaCl, pH 8.0, 10% glycerol, was incubated at room temperature for 1 hour to allow self-binding. It was then diluted to a final concentration of 25nM in either 1X or 4X NuPAGE™ Lithium dodecyl sulfate Sample Buffer (Thermo Fisher) and incubated for 30 minutes at room temperature. Western blotting was performed as described above and membranes were probed with anti-Cx43 (1:1000, Sigma) and anti-Cx43(1:1000, Millipore).

### Protein Modeling

Flexible protein-protein docking was performed using the ATTRACT docking program(de Vries, Schindler et al. 2015). For actin, PDB ID: 2ZWH and PDB ID: 1J6Z were both used in two different runs. For GJA1-20k, PDB ID: 1R5S was used. This PDB file is an ensemble of 10 lowest energy structures of the C-terminus of Connexin 43, all of which were used in each run. HADDOCK(Dominguez, Boelens et al. 2003, van Zundert, Rodrigues et al. 2016) was used to generate harmonic distance restraints in .tbl format to use as parameters for ATTRACT. Ambiguous restraints were generated with R124, P125, R126, P127, L130, E131, and I132 defined as active residues (directly involved in the interaction) and D128 and D129 defined as passive residues (surrounding surface residues). On actin, Y169, a known binding site for RPEL proteins(Mouilleron, Wiezlak et al. 2012), was defined as an active residue. After this information was parsed through ATTRACT, the top 10 models of both actin PDB IDs used were parsed through and models found in most common orientation of GJA1-20k were selected as top models. UCSF Chimera(Pettersen, Goddard et al. 2004) was used for visualization of all structures (color, ribbons, and orientation).

### Computational Alanine Scanning

For the selected models, Robetta Alanine Scanning(Kortemme and Baker 2002, Kortemme, Kim et al. 2004) was used in order to identify hotspot residues on both actin and GJA1-20k in each predicted interaction. Chain 1 was identified as actin, chain 2 was identified as GJA1-20k. Any residue that was identified as having >1ΔΔG(complex) when mutated to alanine is defined as a hotspot residue.

## Acknowledgments

We thank Tara Hitzeman for advice on statistical analysis. We thank Dr. Emil Reisler from the Department of Chemistry and Biochemistry at UCLA for advising us on techniques and the development of this manuscript. This work was funded by R01HL152691, R01 HL138577, R01HL136463, R01HL133286, R01HL136463, and F31HL147404.

## Author contributions

RB, EEG, TTH, and RMS designed research studies. RB, JAP, DS, MW, LK, and SX conducted experiments and acquired data. RB analyzed data. MW conducted computational modeling. QJ and AGK provided patterned coverslips. RB and RMS wrote the manuscript. RB, JAP, MW, DS, QJ, LK, SX, AGK, EEG, TTH, and RMS reviewed and edited the manuscript. TTH and RMS supervised the study.

